# Monkey Facial Dynamics in ‘Minimal Interactions’

**DOI:** 10.1101/246322

**Authors:** Stephen V. Shepherd

## Abstract

Increasingly, researchers are interested in studying the neural substrates of monkey social interactions within laboratory settings. However, semi-naturalistic monkey interactions within lab settings have not been well characterized. We here analyze “minimal interactions” between monkeys in the laboratory. Monkeys, like humans, produce a variety of facial expressions when they encounter one another. It has been unclear what specific information they use to guide their behavior. I recorded the facial signals of captive long-tailed macaques (*Macaca fascicularis*) while they visually interacted while seated in primate chairs. I found that the most consistently-evoked expressions were affiliative in character, most notably in the form of reciprocal lipsmacking. Consistent with prior experiments, lipsmacks were most evident when situational and social ambiguity was maximal: that is, in the first sessions, in the first moments of each session, and when interacting with unfamiliar individuals. Unexpectedly, head and eye orientation played dissociable roles in interactive behaviors. Most intriguingly, monkeys’ facial behaviors reflected both received and previously sent signals, suggesting they interpret others’ current signals in light of their own past communications.

## INTRODUCTION

Many animals live in stable groups with whom they must coordinate their behavior. They do so not only by monitoring others’ actions, but also via the production of communication signals including facial expressions, vocalizations, and body postures. These signals, in turn, influence the probabilities with which subsequent behaviors will be observed and expressed. While admirable work has been done examining sequences of communicative behaviors in the field (Altmann, 1965; Maxim, 1982; Partan, 2002), neuroscientific study of primate social behavior requires bringing it into a laboratory setting. Little is known about sequences of primate social behavior under these conditions.

In nonhuman primates, as in humans, facial movements are crucial communicative gestures. Lipsmacking, for example, is an affiliative signal observed in many genera of nonhuman primates (Hinde & Rowell, 1962; Redican, 1975; Van Hooff, 1962), including apes (Parr, Cohen, & de Waal, 2005). It is characterized by regular cycles of vertical jaw movement, with coordinated lip retraction or puckering. While lipsmacks by both monkeys and apes are often produced during grooming interactions, monkeys (at least) also seem to exchange lipsmacking bouts during more general face-to-face interaction (Ferrari, Paukner, Ionica, & Suomi, 2009; Van Hooff, 1962). I investigated moment to moment changes in lipsmacking and related facial displays across repeated “minimal interactions”.

In the wild, monkeys signal most when interactive outcomes are uncertain (Maxim, 1982). For this reason, I developed a minimal interaction paradigm whereby monkeys (n=8) were abruptly introduced, face-to-face, with another monkey while both were seated in primate chairs; monkeys were paired once with each individual. For this contrived context, I made five predictions. First, I predicted that monkeys should signal most robustly early in each interaction session, because this is when the situational context is least defined. Second, and for the same reason, I predicted monkeys would signal more toward strangers than familiar individuals. Third, I predicted that monkeys would engage in both affiliative (e.g., lipsmacking) and agonistic interactions (e.g., threats and silent bared teeth displays). Fourth, I predicted that monkeys would produce facial signals most robustly when attended, as reflected by either eye or head orientation. Finally, I predicted that lipsmack exchanges should be interdependent.

### MATERIAL AND METHODS

To measure the complexity of monkey signaling behaviors, I developed a “minimal interaction” paradigm. I recorded monkey facial expressions during 5-minute face-to-face interactions. Each interaction was recorded to two digital cameras using a half-silvered mirror, after which each individual’s sequential facial displays were manually scored on a second-to-second basis (Figure 1). Each video showed one interactant; videos were aligned by audio track after scoring. The scorer was blind to the interactive context as well as to the hypotheses of the study. Eight monkeys (male *M. fascicularis*, aged 8-11 years) housed in colonies of four interacted with one another once. This produced 3 interactions in which partners were known from audiovisual contact in their home enclosure and 4 interactions in which partners were relatively unfamiliar. Individual monkeys were involved in no more than one session per day, and all familiar interactions occurred prior to all unfamiliar interactions. All experiments were performed in compliance with the guidelines of the Princeton University Institutional Animal Care and Use Committee.

**Figure 1:**
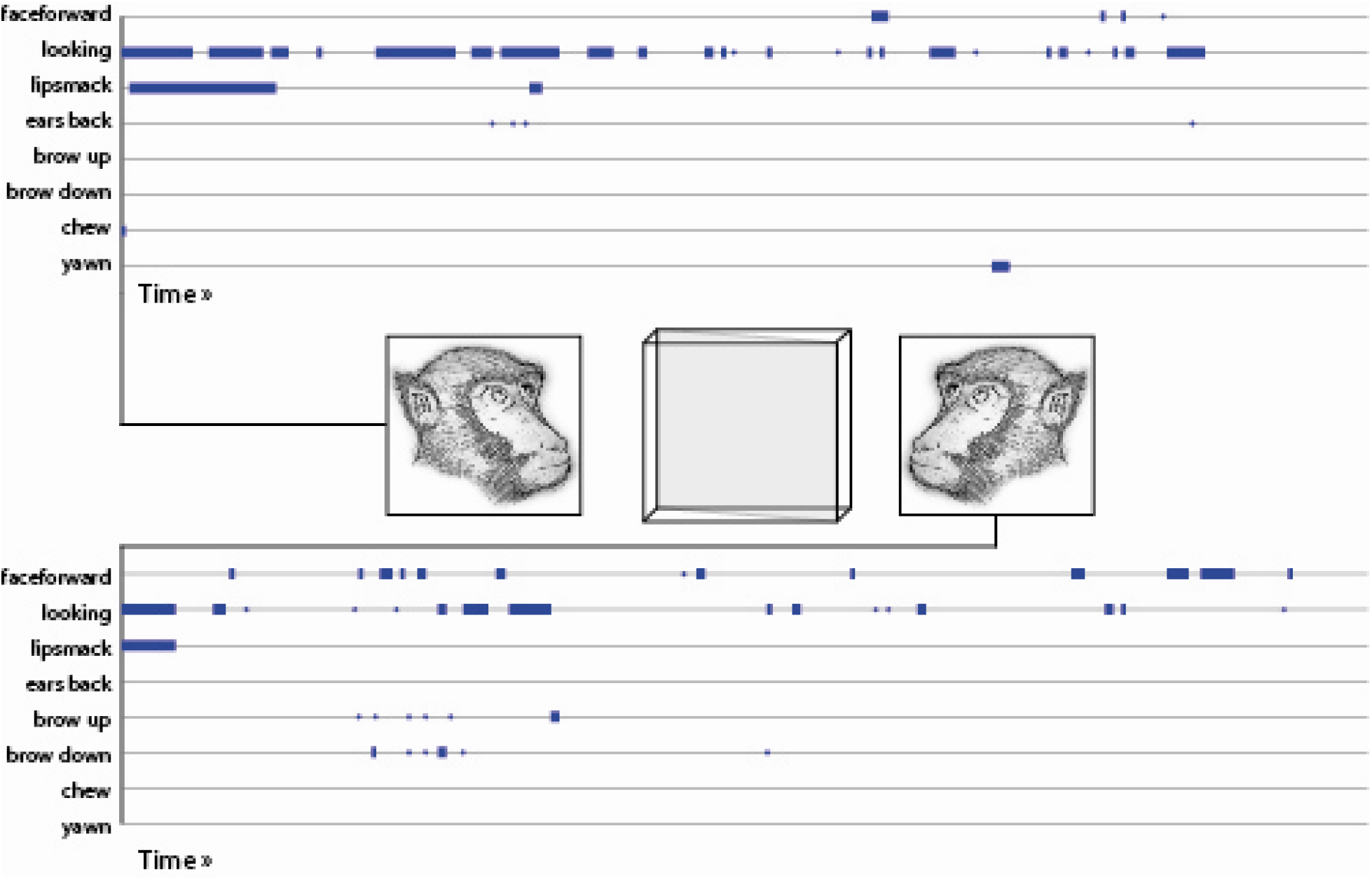
Face-to-face signal exchange. I used a half-silvered mirror to record facial behavior in monkeys interacting face-to-face. Facial movements were scored, and those movements seen in (I) for at least 2 seconds in at least 2 sessions by at least 2 monkeys and (II) for at least 100 seconds total were analyzed in terms of their correlation structure and behavioral consequences.

Fourteen visually-distinct facial postures were scored: facing (head oriented toward partner), looking (eyes oriented toward partner), lipsmacking, brows raised, brows lowered, ears back, ear flick, taut face, head bob, chewing, tongue out, licking, mouth open, and yawn. I analyzed the subset of these which, in the full data set of individual behaviors, were (a) produced for at least 2 seconds in at least 2 interactions by at least 2 monkeys and (b) observed for no less than 100 seconds total across all interactions.

### Comparing signaling early and late in each session

I predicted that monkeys would signal more early in each interaction. To determine how signaling intensity changed over time within sessions, I graphed averages across 10 second bins; significance was determined through Kruskal-Wallis tests (nonparametric ANOVA) over all pooled facial behaviors.

### Comparing signaling between familiar and unfamiliar individuals

I predicted that monkeys would signal most in their earliest interactions, when the experimental paradigm was most novel, and that they would signal more to novel than to familiar individuals. To test this, I examined how total signaling amounts changed across successive interactions. I first examined the significance of session-to-session differences through separate Kruskal-Wallis tests (nonparametric ANOVA) for both “facing” and “looking” orienting responses, and for the pooled average of the six remaining facial behaviors of interest. Based on these results, I tested the revised hypothesis that these behaviors differed between the early sessions (in which they saw familiar individuals) and the later sessions (in which they saw strangers). I first examined the interaction of behavioral type (separately, without pooling) and partner familiarity (familiar or unfamiliar) in predicting total amount of signaling in seconds (ANOVA). FInally, I tested the effect of partner familiarity on each behavioral type individually (Kruskal-Wallis).

### Evaluating the social significance of observed facial behaviors and attention cues

I predicted that both affiliative and aggressive facial displays would be produced during interactions. To determine how different postural features were combined in facial signaling behavior, I measured the Spearman correlation between simultaneously-produced behaviors. I checked the significance of within-subject postural correlations by comparison to a consecutively-preserving time-shuffling permutation baseline: this preserves the discreteness and autocorrelations of each type of behavioral data, while randomizing their interdependence (cf. Shepherd, Steckenfinger, Hasson, & Ghazanfar, 2010).

To evaluate these behaviors’ social function, I correlated facial postures produced in each second with those observed over the preceding 10 seconds using a sliding “boxcar” average. To determine which component of multicomponent signals was most crucial, I considered each observed facial behavior separately while partialing out the effects of the other observed behaviors. I again checked significance through a permutation baseline by randomly pairing individuals’ behavioral sequences, drawn randomly with replacement from all sessions. This served to maintain the time-courses of each behavioral data set while randomizing the dependency between produced and observed signals. This had the additional advantage of controlling for any consistent onset or habituation effects shared across sessions.

### Assessing interdependence between sent and received signals

Finally, to measure the minimal complexity of internal representations guiding monkeys’ signaling behavior, I focused on an important prosocial signal, the lipsmack. Lipsmacks are thought to be reciprocated between monkeys as a means of encouraging association and discouraging aggression or flight. I measured the odds of lipsmacking in any given second as a function of (1) whether the signaler had lipsmacked two seconds prior and (2) whether the target had lipsmacked one second prior. Finally, I compared the change in lipsmack likelihood (i.e. the ratio of odds ratios) seeing a “spontaneous” lipsmack versus seeing a “response” lipsmack. To test significance, I used a permutation baseline composed of simulated interactions, as described above for our between-individual correlation analysis.

## RESULTS

I used a minimal interaction paradigm whereby monkeys (n=8) were abruptly introduced, face-to-face, to another monkey while both were seated; monkeys were paired once with each individual. Eight facial behaviors met our criteria for further analysis: Of the 14 facial behaviors I scored, 11 were observed for more than 2 seconds, in more than 2 sessions, by more than 2 animals. Of 16,800 s total, 3,651 s were spent looking toward the interaction partner, 954 s facing toward the interaction partner, 467 s lipsmacking, 297 s with lowered brow, 212 s with ears flattened, 184 s chewing, 132 s with brows raised, 104 s yawning, as well as 78 s with tensed facial muscles, 72 s licking, and 65 s with the mouth open. The 8 facial behaviors occurring for more than 100 s were subjected to further analysis (Figure 2).

**Figure 2:**
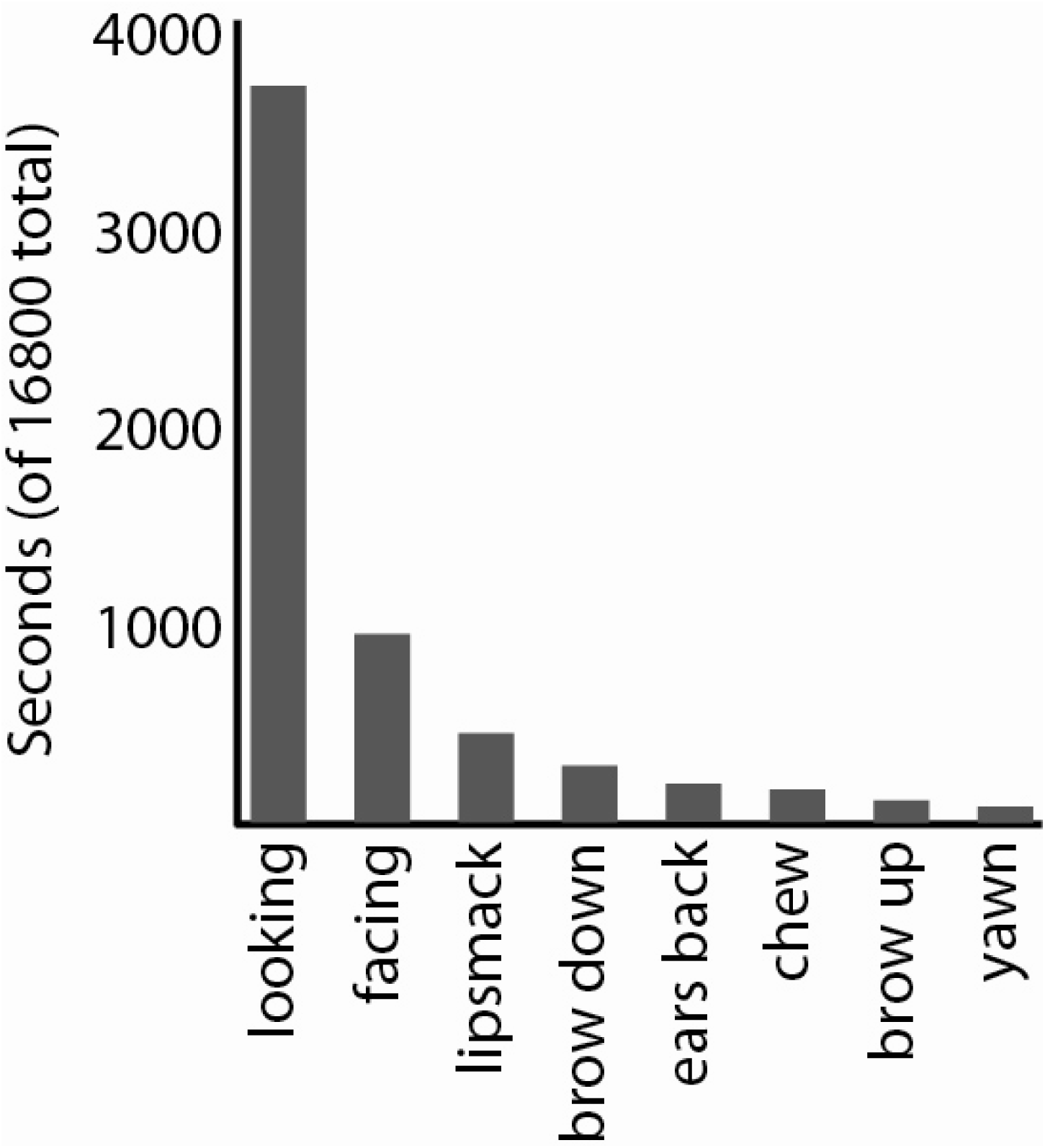
Observed facial behaviors. Of 14 scored behaviors, 11 were produced both for at least two seconds of at least two interactions by at least two individuals, and for over 100 seconds total across all interactions.

In all, I found that monkeys robustly produced facial expressions, most notably the lipsmack gesture. Moreover, monkeys were most socially expressive with strangers and at the beginning of exchanges. I was surprised to find that most interactions in this despotic species were affiliative rather than agonistic, and that two indicators of visual attention—head and eye orientation (facing and looking)—produced different effects on behavior. Finally, I suggest that monkeys’ lipsmack behavior is better predicted by interdependent than independent predictions—they respond differently to individuals to whom they have sent signals, as if they anticipate the consequences their sent signals will have on another’s behavior. I elaborate on each of these findings below.

### Interactions decrease over time within a session

I predicted that monkeys should signal most robustly early in each interaction session, because this is when the situational context is least defined. I found that facial behaviors were indeed most common early in each session (K-S test across all scored behaviors, p=2.4487*10^-190^) (Figure 3). This was especially evident for looking and for lipsmacking, ear flattening, and brow raising, but was not evident for facing or yawning (average time-of-occurrence of facial displays within each 300 s interaction window: facing, 149 s; looking, 118 s; lipsmacking, 46 s; ear flattening, 56 s; brow raising, 69 s; brown lowering, 117 s; chewing, 128 s; yawning, 153 s).

**Figure 3:**
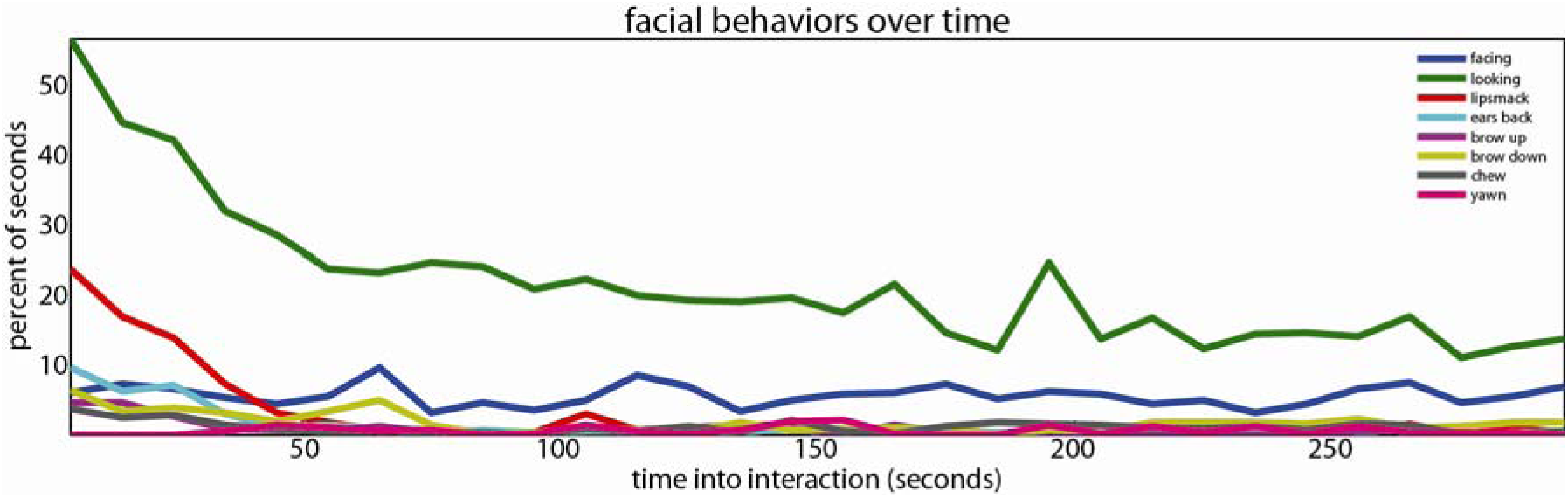
Facial behaviors decreased over time within sessions. Facial behavior likelihood as a percentage of 10-second blocks throughout the interaction window. Most facial behaviors were more likely early in interactions than late, consistent with an attempt to reduce uncertainty about social contexts.

### Increased interactions with unfamiliar versus familiar individuals

For the same reason, I next predicted monkeys would signal more toward strangers than familiar individuals. To look at how signaling evolved between sessions, I broke facial behaviors into the most common types: first, facing; second, looking; and third, all remaining major facial behaviors conducted with mimetic or masticatory muscles (Figure 4). I found that looking and pooled facial expressions changed significantly across sessions (Kruskal-Wallis for looking behavior, Χ^2^(6,49)=15.56, p=0.016; other, Χ^2^(6,49)=18.97, p=0.0042), but facing did not (facing, Χ^2^(6,49)=8.9, p=0.18). While facing tended to decrease across sessions, other facial behaviors showed a stepwise increase after session three, the last session with mutually-familiar participants. This suggested that effects of behavioral uncertainty dominated effects of habituation, but that there were important differences by behavioral type.

**Figure 4:**
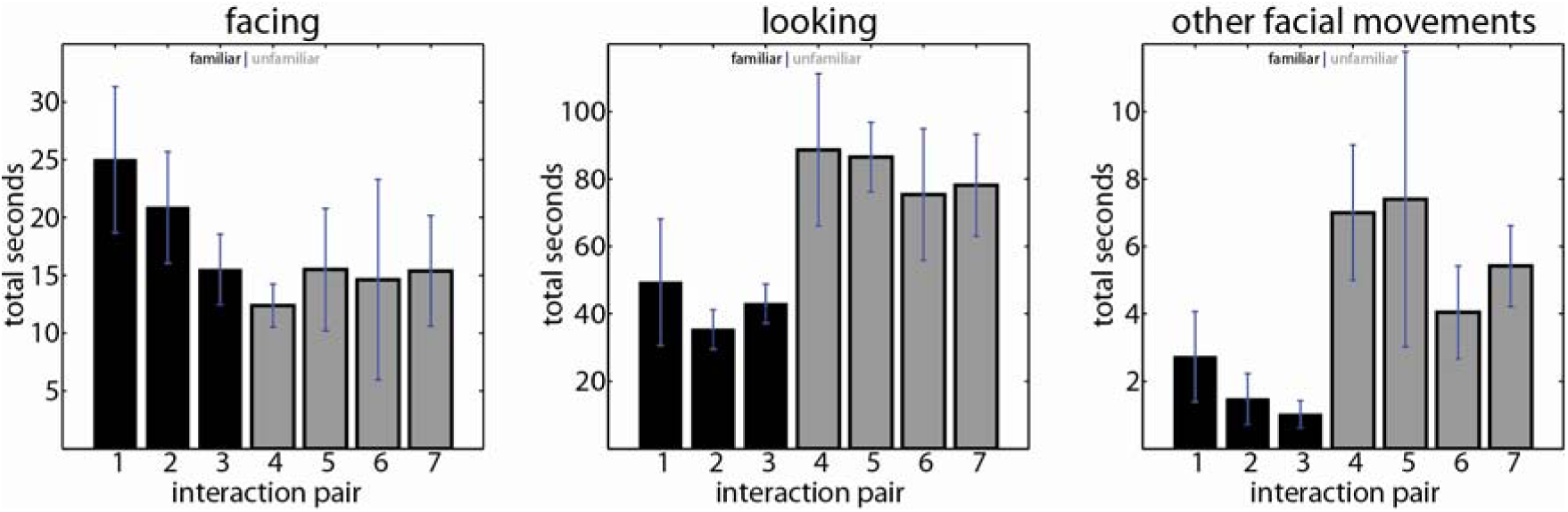
Looking and mimetic facial movement, but not facing behavior, were more common with strangers. Facing behavior decreased across sequential interaction pairs, but looking behavior and non-orienting facial behaviors increased when interacting with unfamiliar partners. These findings are consistent with an attempt to reduce uncertainty about social contexts. As in Figure 2, signaling behavior was most prevalent when behavioral uncertainty was highest.

To explore this further, I considered each behavior type without pooling (ANOVA of seconds signal per session by behavioral type and familiarity, F(1,7,7,432)=9.7, p<10^-10^ for the interaction), then tested each behavior non-parametrically for the influence of familiarity on signal frequency. Most facial behavior was observed at the beginning of interactions and when interacting with strangers, but this varied by behavioral category. Four facial postures significantly increased during the interactions with strangers (Kruskal-Wallis: looking, or eyes-toward-partner, Χ^2^(1,54)=14.26, p=0.00016; lipsmacking, Χ^2^=8.51, p=0.0035; ear flattening, Χ^2^=7.11, p=0.0077; and brow raising, Χ^2^=8.19, p=0.0042). Facing behavior (head toward partner, irrespective of eye direction) was significantly reduced (Kruskal-Wallis, Χ^2^(1,54)=6.19, p=0.013). Brow lowering also trended more common when viewing strangers, while chewing and yawning trended less common.

### “Minimal interactions” produce robust affiliative signaling

I predicted that monkeys would engage in both affiliative (e.g., lipsmacking) and agonistic interactions (e.g., threat or submission displays). Contrary to this latter prediction, I observed no outright open mouth threat or silent bared teeth displays. To further investigate, I examined the co-occurrence of facial behaviors both within and between signalers (Figure 5). A cluster of mutually-associated behaviors (permutation test, p<0.05) were produced by individuals: looking was correlated with every other facial behavior, and lipsmacking, ears back, and brow up were mutually correlated. By contrast, yawns were produced less often than chance in the same second as lipsmacks / flattened ears / raised brows (likely due to simple biomechanical incompatibility).

**Figure 5:**
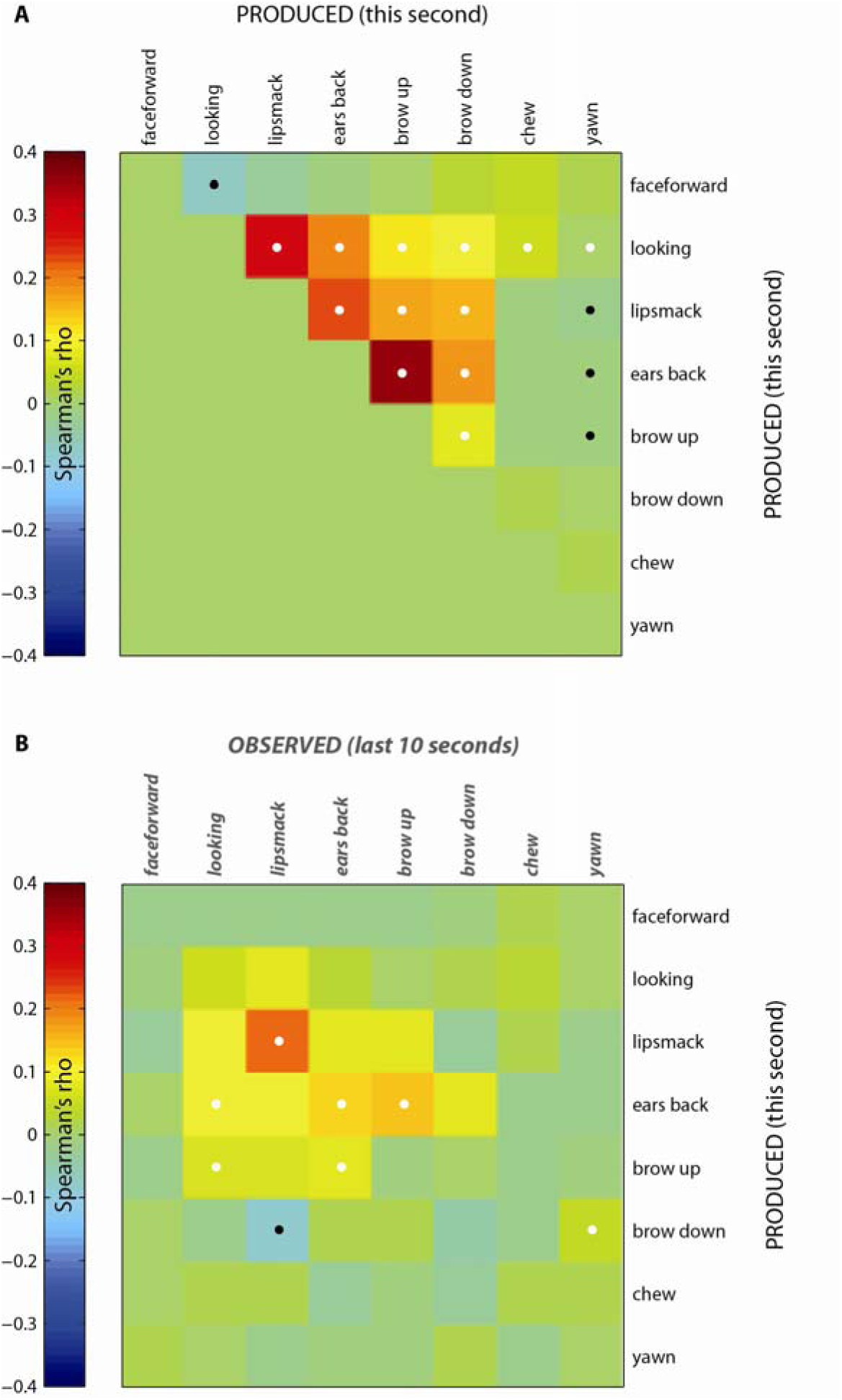
Face-to-face signal correlogram. (**A**) Simultaneously produced signals (Spearman’s rho) suggest a cluster of related looking-and-interaction behaviors, contrasted with facing behaviors and yawning. (**B**) Partial correlations between signals produced (rows) and received over the last 10 seconds (columns; partial correlations distinguish effects of each specific received signal type) indicate affiliative signals (looking, lipsmacking, ears flattening, and brow raising) were not only co-produced but were also matched by partners. This sort of behavioral mimicry has been suggested by prior field and lab studies, and suggests that like humans, monkeys may reflexively reciprocate prosocial greetings. In both panels, white dots mark squares which were significantly higher than a permutation baseline, while black dots mark squares which were significantly lower. Permutation was performed by shuffling time, for self-correlations in **A**, and by shuffling trial number, for inter-individual correlations in **B** (α p=0.05, two-tailed).

To assess the causal influence of signals on observers, I examined how produced behaviors were correlated with the average amount of observed behaviors over the prior 10 seconds. I used partial correlation to distinguish which components of any multicomponent displays (e.g. lipsmack / flattened ears / raised brows) were most strongly causal connected. I found significant patterns of causal influence (p<0.05 by permutation):

- Lipsmacks elicited lipsmacks (r=0.22) and suppressed brow lowering (r=0.08).
- Flattened ears and raised brows were produced more when looked toward, or when partners flattened their ears. Looking elicited both ear flattening (r=0.11) and brow raising (r= 0.07); ear flattening likewise elicited ear flattening (r=0.13) and brow raising (r=0.08). Brow raising elicited ear flattening (r=0.14).
- Yawning elicited lowered brows (r=0.04).

The most commonly observed pattern in minimal interactions is thus the reciprocal exchange of the multicomponent lipsmack display, which includes both obligate patterned orofacial movement in the perioral mimetic muscles and non-obligate brow raising and ear flattening (Shepherd, Lanzilotto, & Ghazanfar, 2012). This was a partial confirmation of our initial hypothesis that both affiliative and aggressive signals would be elicited in a “minimal interaction” paradigm: affiliative signals were observed, but I saw little or no agonistic signaling, including a total absence of their canonical expressions (open-mouthed threats, signaling dominance, or silent bared-teeth, signaling subordination) and only fleeting appearances of stress- and conflict-related signals like yawning and brow lowering.

### Head and eye orientation play different roles in communication

Our fourth prediction was that monkeys would produce facial signals more robustly when attended, as assessed through observed eye and head orientation (Figure 6, cf. Figure 5). However, I discovered that facing behavior and looking behavior followed very different time courses within each interaction session, and responded differently to familiarity. Moreover, on a second-to-second basis, facing was anticorrelated with looking—perhaps to avoid staring, an agonistic signal in macaques. I examine the relationship of attention cues to co-produced facial behaviors (“produced with”), to observed facial behaviors which preceded instances of the facing/looking behavior (“elicited by”), and to observed facial behaviors which followed instances of facing/looking behavior (“provoked”). I found that facing and looking behavior played strikingly different roles in social exchange, rather than comprising a single signal of directed attention as generally assumed (e.g. in the context of gaze-following behavior; cf. Shepherd, 2010). Specifically, facing and looking were correlated with different behaviors on a second-by-second basis and were themselves anticorrelated (permutation test, p<0.05; facing was significantly less correlated than looking with lipsmacking, ear flattening, brow raising, and brow lowering).

**Figure 6:**
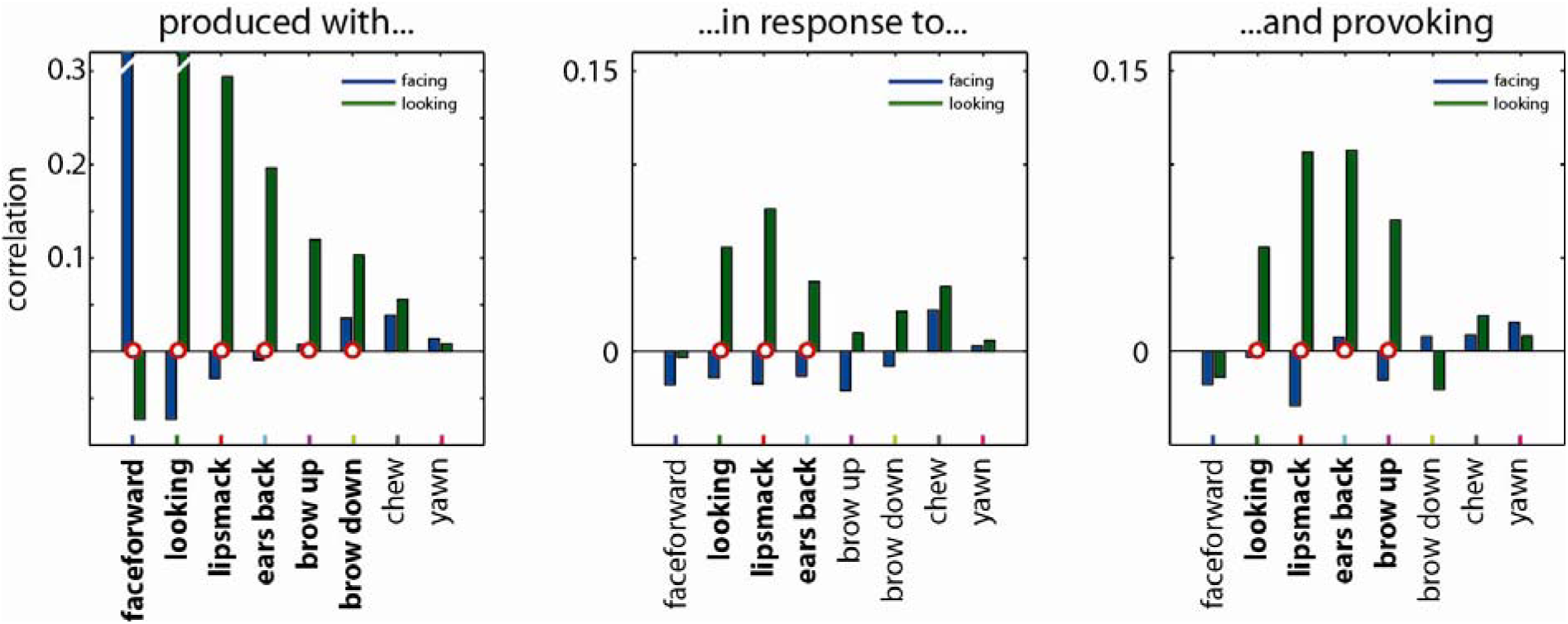
Head and eye orientation cues play different roles in dyadic behavior. Though head and eye direction both imply direction of attention (Shepherd 2010), they play strikingly different roles in dyadic social behavior. Facing behavior was weakly predictive of co-produced facial behaviors and of social responses, and was anticorrelated with gazing (perhaps to avoid producing aggressive staring displays). Looking was correlated with affiliative signal production in these “minimal interactions” for both the subject and the respondent.

Moreover, looking was more often than facing produced in response to looking, lipsmacking, and ear flattening (p<0.05); it was similarly more likely than facing to elicit looking, lipsmacking, ear flattening, and brow raising (p<0.05).

### Lipsmack exchanges modeled as dialogs

Finally, I quantitatively addressed the minimal complexity required to explain animals’ lipsmacking behavior: Is signal exchange determined solely by the present state of both actors, or by their interaction history? To illustrate, consider the human exchange of smiles. Let’s say one wanted to predict whether an individual, Ann, is smiling towards Bill. Both Ann and Bill’s past behavior will influence Ann’s decision. Was Ann smiling towards Bill a second ago? Did Bill smile towards Ann? These data should not be considered wholly independently. The important thing is that each responds appropriately to the other’s greeting. Has anyone smiled? If Ann smiled toward Bill, has he replied? If Bill smiled towards Ann, has she? The crucial point is that this inter-dependence between sent and received signals is a hallmark of human social behavior and is mathematically detectable. To quantitatively address the complexity of animal social interactions, I can determine whether the behavior of interactants is better predicted by *interdependent* versus *independent* signal histories.

I therefore asked whether monkeys responded differently to lipsmacks observed in different interactive contexts: Do monkeys distinguish spontaneous signals from those that are mere replies? I found that like humans, monkeys represent communication signals in terms of their dyadic context. Specifically, I found that the monkeys were significantly less likely than baseline to lipsmack when they hadn’t received a lipsmacked (permutation baseline, p<0.05, both when subject had or hadn’t lipsmacked two seconds prior), but significantly more likely than baseline to lipsmack to an unsolicited lipsmack (p<0.01) (see Table 1 and Figure 7). In total, lipsmack production odds were 6.3 times greater after seeing an unsolicited lipsmack then after seeing a lipsmack that had been previously solicited or acknowledged, though this measurement did not attain statistical significance (permutation baseline, p>0.05). These data suggest that monkeys differentiated between initiating and response lipsmacks, weighing the former much more heavily than the latter.

**Table 1:**

| LIPSMACK LIKELIHOOD | Unreceived <sub>1 second ago</sub> | Received <sub>1 second ago</sub> |
| --- | --- | --- |
| Unsent <sub>2 seconds prior</sub> | **0.6%** | **9.8%** |
| Sent 2 seconds prior | 64% | 83% |

**Figure 7:**
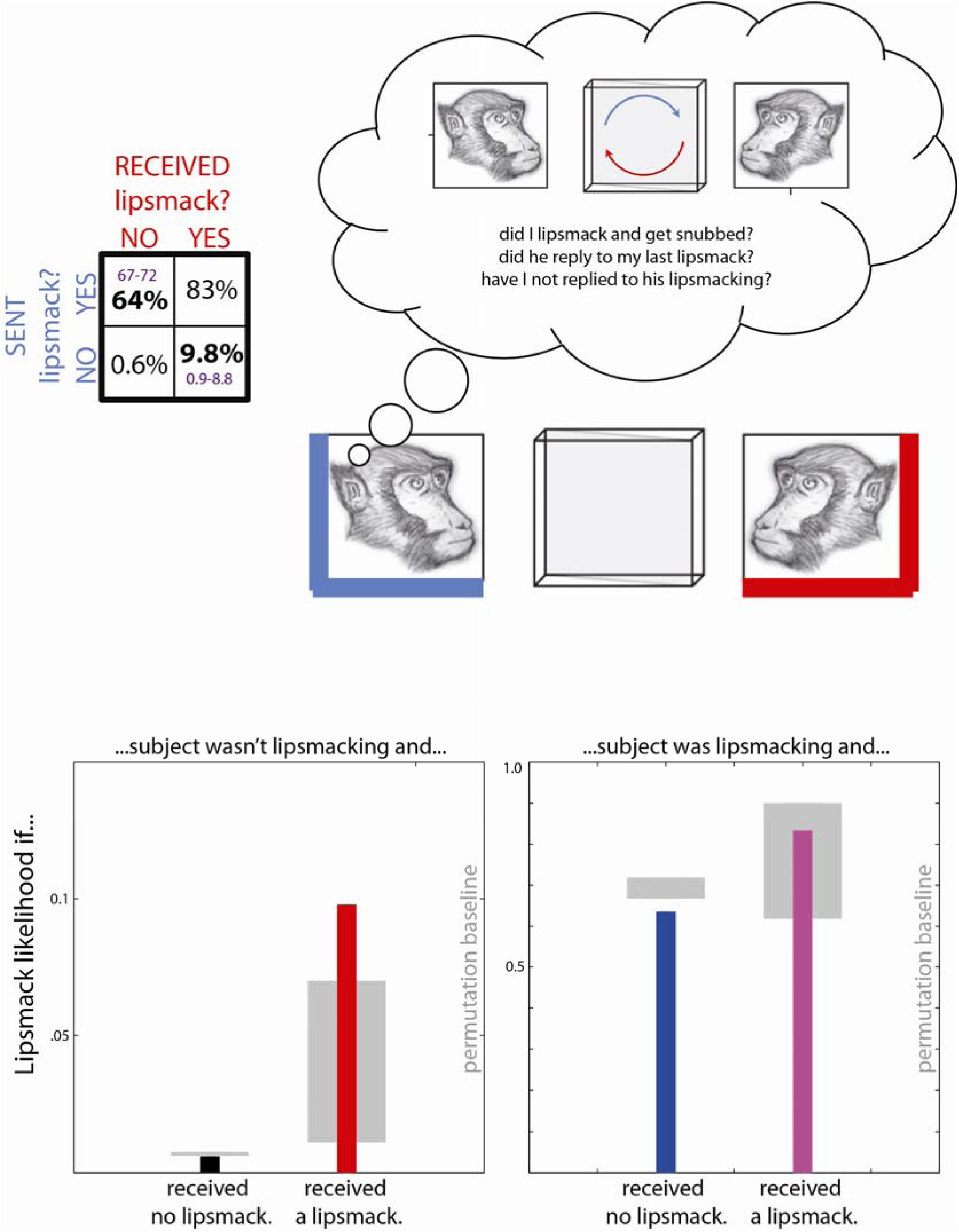
Lipsmack exchange as a dialog. Patterns of sent and received lipsmacks were not independent but interdependent: Given the rate of lipsmacking observed in the experiment, lipsmacking was more likely than chance when responding to “spontaneous” partner lipsmacks, and less likely than chance when produced lipsmacks were “snubbed”. These “spontaneous” partner lipsmacks has an 6 times greater impact on odds of lipsmack production than did “responsive” partner lipsmacks.

## DISCUSSION

Of 14 facial behaviors scored, 8 were found to arise repeatedly in these “minimal” interactions and to total at least 100 seconds duration, our threshold for further analysis. These behaviors included orienting movements (facing, looking), movements of facial muscles (lipsmacking, brow lowering), and movements of jaw muscles (yawns, chews). The most vigorous facial behavior arose at moments of increased uncertainty – early in interactions, and more generally when meeting strangers. The finding that many signals increased when meeting unfamiliar monkeys is consistent with prior literature, which has suggested these signals reduce behavioral uncertainty (Maxim, 1982) and so reduce the risk of conflict. Interestingly, movements of the facial muscles increased when viewing strangers, while movements of jaw muscles did not, consistent with reports that the former have been optimized for communicative behavior (Dobson, 2009; Dobson & Sherwood, 2011; Shepherd et al., 2012).

Observed behaviors included a suite of related displays: lipsmacking, brow raising and ear flattening regularly co-occurred and elicited similar responses in viewers, most notably including reciprocal lipsmacking. Reciprocation of lipsmacking has been observed both in the wild (Andrew, 1963; van Hooff, 1967) and in captivity (Livneh, Resnik, Shohat, & Paz, 2012; Mosher, Zimmerman, & Gothard, 2011). I replicate the finding that lipsmacking, brow raising and ear flattening are behaviorally associated, but do not see evidence that they are strictly synonymous. This may suggest they carry distinct but related meanings, consistent with our finding that the fine dynamics of eyebrow raising and ear flattening are decoupled from those of lipsmacking (Shepherd et al., 2012).

The robust reciprocity of affiliative lipsmacking in monkeys is interesting by way of comparison to humans. There is a large body of work on similar prosocial mimicry in humans: Humans reflexively mimic facial expressions (Dimberg, Thunberg, & Elmehed, 2000) and mimic behavior more when more motivated to be liked (Lakin & Chartrand, 2003); furthermore, such matching behavior is successful in increasing liking (Chartrand & Bargh, 1999). If lipsmacking, like gaze (Shepherd, 2010), is pervasively and reflexively mimicked, then this behavior will be among the most elementary of social sensorimotor behaviors.

However, just as these social behaviors can be simple and reflexive, they can also have hidden complexity (e.g. Shepherd, Deaner, & Platt, 2006). I found that not all lipsmacks are equal: “Response” lipsmacks are less effective than “spontaneous” lipsmacks at inducing lipsmacking behavior. This result suggests that monkeys who have recently lipsmacks are refractory to observed lipsmack signals. Monkeys may recognize lipsmack exchange as a kind of dialog, in which received signals are interpreted in light of previously sent displays. Our data are likewise consistent with goal-oriented behavior, in which lipsmacks are used instrumentally to influence another’s behavioral state. Prior research had suggested that remembered interaction history is crucial in organizing responses to vocalizations (Cheney & Seyfarth, 1997; Engh, Hoffmeier, Cheney, & Seyfarth, 2006; Wittig, Crockford, Seyfarth, & Cheney, 2007), likely because the targets of an auditory signal are underspecified by readily-observable environmental cues. Here, I show that immediate interactive context is similarly crucial to the interpretation and governance of visual signal production in dyadic interactions, in which signal targets are certain but intent may be ambiguous.

Facial expressions play a major role in close-range primate interactions (Altmann, 1965; Partan, 2002) and their effective use appears to be crucial in navigating large social groups ((Dobson, 2009), see also (Parr, Waller, & Fugate, 2005)). For facial expressions to optimally coordinate behavioral states, however, monkeys should interpret received signals in light of those they have already sent. Our data indicate that they do this: Monkeys, like humans, understand signals in terms of their interaction context. I hypothesize that specific neurons are sensitive to relational features of signal exchange, with nonlinear sensitivity to combinations of previously-sent and recently-received signals. Such sensitivity could in principle take several forms, not all of which require sophisticated cognitive mechanisms: Sent signals may reflect internal states that change either perceptual processing or expressive thresholds, or may explicitly influence representations of the external world through predicted consequences for the behavioral predispositions or mental states of others. These possibilities will be most efficiently disambiguated through further behavioral research and, especially, through neurophysiological investigation.

## ACKNOWLEDGEMENTS

I am indebted to Matthew Slayton for video coding of monkey facial expressions and to Daniel Takahashi for technical suggestions. This work was conducted in the laboratory of Asif A. Ghazanfar at Princeton University; I am grateful for both his advice and for his patience when I failed to heed it.

